# *Drosophila* storage proteins promote both the rate and the duration of tumor growth

**DOI:** 10.1101/2025.10.02.680002

**Authors:** Luca Valzania, Dalmiro Blanco-Obregon, Aya Alami, Pierre Léopold

## Abstract

Amino acid storage proteins, such as serum albumin in mammals, are known to accumulate within tumors, but their precise contribution to tumor biology remains unclear. Using both cachectic and non-cachectic tumor models in *Drosophila*, we observed that fat body-derived hexamerins accumulate in the tumors through a selective uptake process. Disabling this uptake led to a marked reduction in tumor growth, demonstrating that hexamerins are used as a nutrient source to support cancer progression. Hexamerin uptake also supports expression of the relaxin-like Dilp8 by the tumor, inhibiting ecdysone production and extending the growth period. This coupling between nutrient uptake by the tumor and inhibition of the developmental progression exerts a full diversion of the host resources. Functional parallels with mammalian albumins suggest evolutionarily conserved mechanisms with potential implications for cancer biology.

## INTRODUCTION

A fundamental challenge faced by rapidly growing tumors is to secure sufficient nutrients to support their relentless anabolic demands. Since Warburg’s early recognition of altered carbohydrate metabolism in cancer cells(*1*), a wide range of low molecular weight nutrients, including fatty acids, ketone bodies and amino acids, have been investigated as critical contributors to tumor growth and survival. Yet despite decades of study, a comprehensive understanding of tumor nutrient metabolism and energy requirements remains elusive(*2*). As a result, identifying additional nutrient sources that support tumor proliferation continues to be a major focus in cancer research.

One historically underexplored nutrient source is plasma proteins. As early as 1948, Mider and colleagues suggested that plasma proteins such as serum albumin might serve as nitrogen donors for tumors(*3*). Subsequent studies confirmed that albumin can accumulate in the tumor microenvironment and be taken up by cancer cells(*4*). Nevertheless, the concept of plasma protein catabolism as a direct amino acid supply for proliferating tumor cells has not been widely investigated.

In parallel, the use of albumin as a drug carrier in cancer therapy has gained significant attention over the past two decades, highlighting its biological importance in tumors(*5*). Despite this interest, the molecular players mediating albumin uptake, such as specific receptors and binding proteins, remain poorly characterized(*6*). In mammals, a major obstacle to investigating albumin’s role in tumor biology is its constitutive synthesis throughout life. By contrast, insects offer a tractable model to study storage proteins with precise temporal resolution. In these organisms, functional homologs of albumin are represented by hexamerins, which share multiple key features with mammalian serum albumin. They are the most abundant proteins in the circulatory system, they are primarily synthesized by an organ analogous to the liver, they act as dynamic reservoirs of amino acids, they support metabolic demands throughout the life cycle, they bind and transport small hydrophobic molecules and hormones, and their synthesis is tightly regulated by hormonal and nutritional cues(*7*). Due to this ensemble of functional similarities, hexamerins provide a unique opportunity to investigate how storage proteins contribute to tumor metabolism. In *Drosophila melanogaster*, hexamerins comprise the Larval Serum Proteins 1 and 2 (LSP1, LSP2) and the Fat Body Proteins 1 and 2 (FBP1, FBP2)(*7*). These storage proteins are synthesized by the fat body during a defined window of larval development and accumulate in the hemolymph. Subsequently, LSPs and FBP2 are re-imported into the fat body via FBP1, where they serve as essential nutrient reserves for metamorphosis(*8*).

*Drosophila* offers powerful genetics to manipulate both tumor cells and host tissues, enabling a detailed, tissue-specific analysis of nutrient exchanges. While prior studies have revealed tumor-induced metabolic rewiring and systemic wasting reminiscent of cancer cachexia in flies(*9*), the role of macromolecular nutrients such as hexamerins has not been explored. In this study, we identify a mechanism by which epithelial tumors in *Drosophila* larvae hijack hexamerin resources to promote their own growth. We show that tumors actively internalize hexamerins through FBP1, and that loss of FBP1 impairs hexamerin uptake and limits tumor expansion. In addition to nutrient hijacking, tumors exert profound systemic effects on their hosts, including developmental delay, metabolic dysfunction, and cachexia-like wasting. In *Drosophila*, epithelial tumors can secrete the relaxin-like hormone Dilp8, which coordinates tissue growth and delays metamorphosis in response to local stress or growth perturbations(*10–14*). Here, we demonstrate that hexamerin uptake by tumors is required to induce *dilp8* expression and delay pupariation, thereby providing tumors with extended time for growth.

Using *Drosophila* to dissect nutrient fluxes between the host and the tumor, our work uncovers a tumor-driven strategy to best exploit host storage proteins and the developmental time window for growth.

## RESULTS

### Fat body-derived hexamerins accumulate in Yki^S168A^ tumors

To investigate the role of amino acid storage proteins in a tumor context, we used a *Drosophila* model of epithelial tumorigenesis based on the tissue-specific overexpression of a constitutively active form of Yorkie (*UAS-yki*^*S168A*^*-gfp*), the fly homolog of the mammalian oncogene YAP(*15, 16*). Expression was driven in the wing imaginal disc using the *pdm2-Gal4* driver. This manipulation overrides Hippo pathway inhibition, resulting in uncontrolled tissue growth and a failure to initiate pupariation (Fig. S1A). As a consequence, tumor-bearing larvae enter an extended larval stage lasting ∼ 8 days and ultimately die without forming pupae (Fig. S1A). These tumors, which initiate during the third larval instar, expand significantly and reach volumes up to 15 times larger than those of *wild-type* wing discs(*17*). Notably, this developmental window coincides with the peak of hexamerin production(*8*), making this model well-suited for studying tumor/albumin-like protein interactions.

We started asking whether these tumors accumulate hexamerins, similar to the accumulation of albumin and other nutrient carriers observed in some mammalian cancers. To this end, we used custom-generated antibodies against LSPs and FBPs(*8*) to perform immunofluorescence staining on *pdm2>yki*^*S168A*^*-gfp* tumor tissue. Although hexamerin presence in the wing imaginal discs is typically very low (Fig. 1A), we observed a striking accumulation of both LSPs and FBPs in tumor-bearing discs (Fig. 1B, upper row). The signal was particularly intense at tissue folds, raising the possibility of non-specific trapping of the antibodies. However, high-resolution confocal microscopy revealed that hexamerins were localized in vesicular structures within the cytoplasm of tumor cells (Fig. 1B, lower row), commonly viewed as a sign of intracellular uptake.

**Fig. 1.**
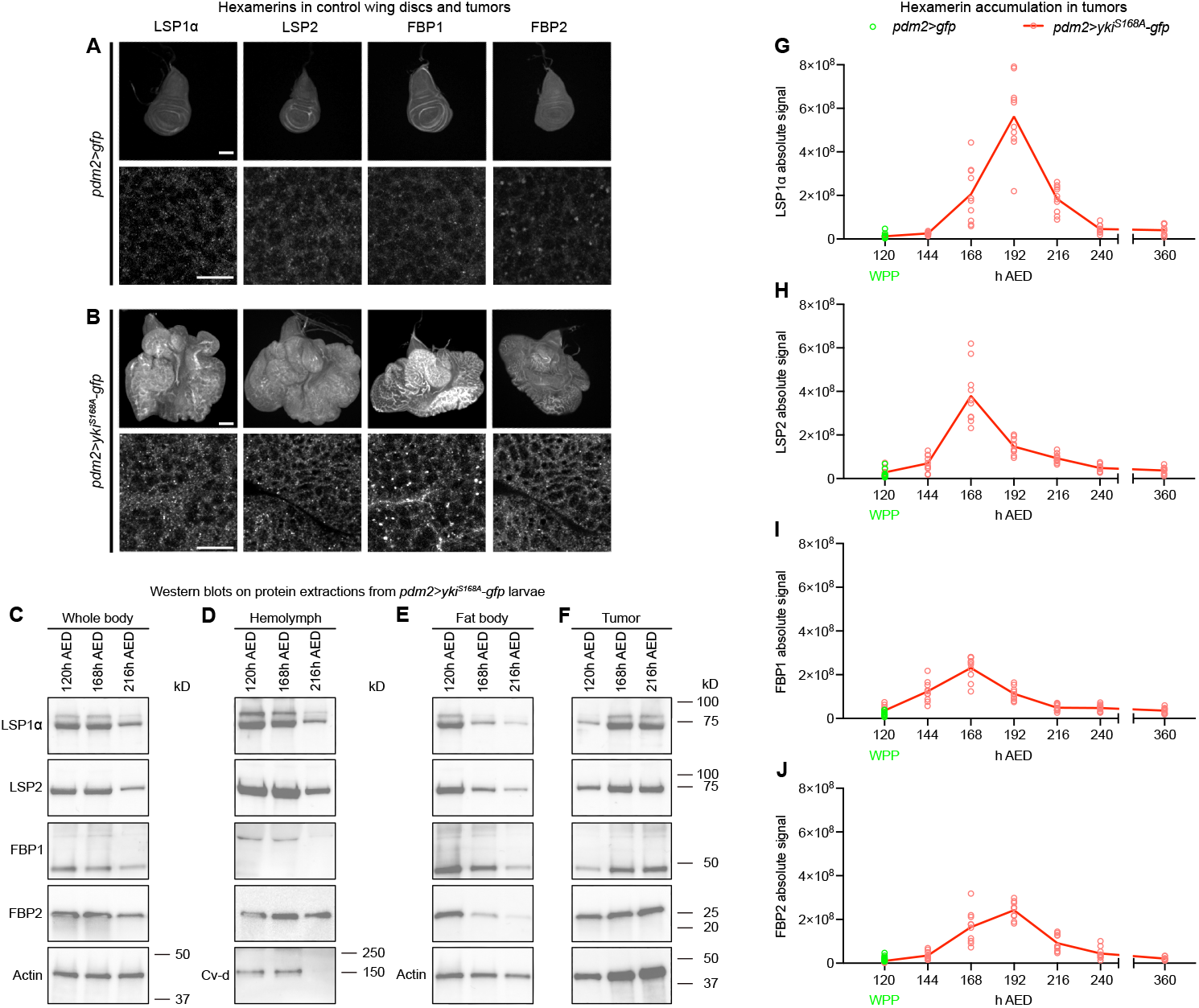
Yki^S168A^ tumors sequester circulating hexamerins, diverting them from normal reuptake by the fat body. (**A**) Representative wing imaginal discs dissected at 120h after egg deposition (AED) from *pdm2>gfp* white prepupae, stained with antibodies against LSP1α, LSP2, FBP1, and FBP2. Top row: entire discs shown; scale bar = 100μm. Bottom row: high-magnification views of the wing pouch show hexamerins localized inside disc epithelial cells; scale bar = 10μm. (**B**) Representative tumors from wing discs dissected from *pdm2>yki*^*S168A*^*-gfp* larvae stained with the same antibodies. Top row: full tumor views; scale bar = 100μm. Bottom row: high-magnification images show robust hexamerin accumulation within tumor cells; scale bar = 10μm. (**C-F**) Immunoblots of protein extracts from whole body (**C**), hemolymph (**D**), fat body (**E**), and tumors (**F**) of *pdm2>yki*^*S168A*^*-gfp* larvae collected at indicated time points (h AED). Blots were probed with antibodies against LSP1α, LSP2, FBP1, FBP2, and loading controls Actin and Cv-d. Molecular weight markers are indicated on the right. (**G-J**) Quantification of LSP1α (**G**), LSP2 (**H**), FBP1 (**I**), and FBP2 (**J**) levels through immunostaining in tumors during the extended third instar stage of *pdm2>yki*^*S168A*^*-gfp* larvae. Timepoints (in h AED) are shown on the x-axis. The 120h AED timepoint corresponds to the pupariation of *pdm2>gfp* control larvae, from which hexamerin levels were quantified in wing disc pouch. For each timepoint, 10 individual discs or tumors were analyzed.

To determine the origin of the hexamerins accumulating in tumors, we first compared the mRNA levels of all hexamerin-encoding genes in wing imaginal discs and fat body tissue. As expected, expression of these genes was high in the fat body and very low in *wild-type* wing discs (Fig. S1B), consistent with the fat body being the primary site of hexamerin synthesis(*8*). In wing discs from tumor-bearing larvae, expression was even further reduced, ruling out local production by the tumor as the source of hexamerin accumulation (Fig. S1C). We next analyzed hexamerin gene expression in fat body tissue from control (*pdm2>gfp*) and tumor-bearing (*pdm2>yki*^*S168A*^*-gfp*) animals to assess whether hexamerin production is modified by the presence of the tumors. We found no significant changes in expression of *Lsp1* (Fig. S1D, E), *Lsp2* (Fig. S1F, G) or *Fbp2* (Fig. S1J, K) between the two conditions. The only notable difference was a reduction in *Fbp1* expression in *pdm2>yki*^*S168A*^*-gfp* larvae at 128h after egg deposition (AED) (Fig. S1H, I). This is consistent with a reduced production of ecdysone, the major steroid hormone regulating insect development, in these animals(*18–20*) and with the fact that *Fbp1* expression relies in part on ecdysone(*8*).

We next investigated how hexamerin distribution changes during tumor progression. To do this, we performed Western blot analyses on protein extracts from whole larvae, hemolymph, dissected fat body, and tumor tissue at multiple time points throughout the extended larval stage (for details see Materials and Methods). In tumor-bearing animals, total hexamerin levels in whole-body extracts decreased progressively over time (Fig. 1C, see normalization to actin levels in Fig. S1L). A similar decrease was observed in the hemolymph (Fig. 1D, Fig. S1M: hemolymph levels are normalized to Cv-d(*21*), however, this lipoprotein is no longer detectable at 216h AED precluding normalization for this time point) and in the fat body (Fig. 1E, Fig. S1N). By contrast, we observed a progressive increase of hexamerin signals in the tumor paralleled with an increase in actin levels, reflecting the presence of elevated hexamerin levels during the first phase of tumor growth (Fig. 1F, Fig. S1O).

To further evaluate the dynamics of hexamerin levels in the tumor, we developed a custom Fiji macro (see Materials and Methods and the code in Fig. S2) to measure hexamerin immunofluorescence from full confocal z-stacks. The integration of the fluorescent signal over entire tumors revealed that hexamerin levels increase during the first two to three days of the extended larval period, then start declining (Fig. 1G-J).

Collectively, these findings indicate that hexamerins are produced by the fat body, secreted into the hemolymph and subsequently taken up by tumor cells.

### FBP1-dependent uptake of hexamerins enables tumor growth

We next sought to determine the mechanism by which hexamerins are taken up by tumors. In the normal developmental context, hexamerins re-enter the fat body only when bound to a specific serum protein called FBP1, which is produced by fat cells at the onset of metamorphosis(*8*). This raised the question of whether FBP1 is also required for hexamerin uptake by tumors.

To address this, our initial strategy was to use two separate ectopic expression systems: one to induce tumors in the wing disc, and a second to knockdown *Fbp1* in the fat body. However, due to technical constraints, we instead employed two Gal4/UAS system to simultaneously induce Yki^S168A^-driven tumor growth in the wing disc and silence *Fbp1* expression in the fat body. Because this approach results in ectopic Yki activation in the fat body and *Fbp1* knockdown in the wing disc, we performed a set of control experiments to rule out unintended developmental effects that could confound our interpretation. Specifically, we assessed larval feeding behavior during the third instar (Fig. S3A), wing disc volume (Fig. S3B) and hexamerin expression at 120h AED (Fig. S3C-H), timing of pupariation (Fig. S3I) and eclosion (Fig. S3J), and adult wing size (Fig. S3K) across all relevant genotypes. These analyses did not reveal phenotypes caused by ectopic Yki activation in the fat body or *Fbp1* knockdown in the tumor, except for a slight delay at pupal eclosion and a mild reduction of adult wing size (Fig. S3J, K) as previously reported for *Fbp1* depletion in the fat body(*8*).

We then performed immunofluorescence quantification of hexamerins in tumors in the presence or absence of FBP1. Notably, *Fbp1* expression, already at very low levels in the wing disc, further declined over time in tumor-bearing discs (Fig. S4A), suggesting that hexamerin uptake in the tumor is not driven by local *Fbp1* expression. While in control tumor-bearing larvae, hexamerins robustly accumulated in tumor tissues (Fig. S4B-E), they were barely detectable upon *Fbp1* knock-down (Fig. S4B-E). Western blot analysis on hemolymph samples from *UAS-Fbp1*^*RNAi*^*/UAS-yki*^*S168A*^*-gfp;pdm2-Gal4/Lpp-Gal4* larvae showed sustained storage protein levels (Fig. 2A, to compare with Fig. 1D), suggesting that FBP1 depletion, by preventing hexamerin uptake both in fat body and in tumor cells, induces their accumulation in the hemolymph.

**Fig. 2.**
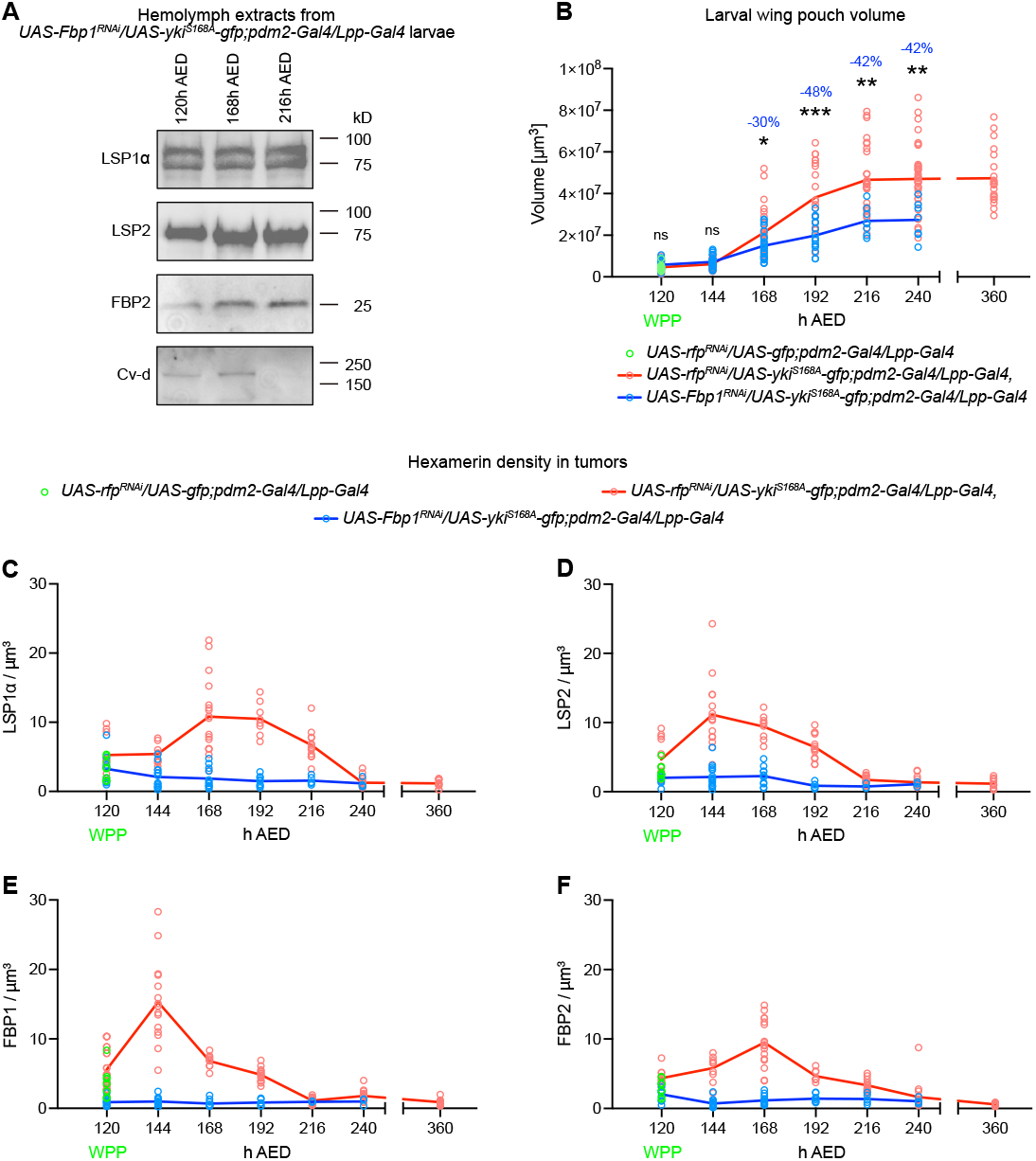
Hexamerins enter Yki^S168A^ tumors via a FBP1-dependent mechanism and support tumor growth. (**A**) Immunoblots of hemolymph extracts from *UAS-Fbp1*^*RNAi*^*/UAS-yki*^*S168A*^*-gfp;pdm2-Gal4/Lpp-Gal4* larvae collected at the indicated time points (h AED). Blots were probed with antibodies against LSP1α, LSP2, FBP2, and Cv-d (loading control). Molecular weight markers are shown to the right. (**B**) Quantification of wing pouch volume (y-axis) in tumor-bearing larvae, either in the presence (*UAS-rfp*^*RNAi*^*/UAS-yki*^*S168A*^*-gfp;pdm2-Gal4/Lpp-Gal4*) or absence (*UAS-Fbp1*^*RNAi*^*/UAS-yki*^*S168A*^*-gfp;pdm2-Gal4/Lpp-Gal4*) of FBP1. Volumes were measured at the indicated time points (x-axis, in h AED). Wing pouch volume of control larvae (*UAS-rfp*^*RNAi*^*/UAS-gfp;pdm2-Gal4/Lpp-Gal4*) was analyzed at pupariation (120h AED). A total of 7 to 38 wing pouches were analyzed per genotype at each time point. Statistical significance was assessed using unpaired *t*-tests or one-way ANOVA, as appropriate. ns = not significant; * = *p* ≤ 0.05; ** = *p* ≤ 0.01; *** = *p* ≤ 0.001. The percentage reduction in wing pouch volume in the FBP1-depleted condition (*UAS-Fbp1*^*RNAi*^*/UAS-yki*^*S168A*^*-gfp;pdm2-Gal4/Lpp-Gal4*) relative to *UAS-rfp*^*RNAi*^*/UAS-yki*^*S168A*^*-gfp;pdm2-Gal4/Lpp-Gal4* between 168h and 240h AED is indicated above the asterisks. No surviving *UAS-Fbp1*^*RNAi*^*/UAS-yki*^*S168A*^*-gfp;pdm2-Gal4/Lpp-Gal4* larvae were observed at 360h AED. (**C-F**) Quantification of intratumoral density of LSP1α (**C**), LSP2 (**D**), FBP1 (**E**), and FBP2 (**F**) during the prolonged third instar stage in larvae of the following genotypes: *UAS-rfp*^*RNAi*^*/UAS-yki*^*S168A*^*- gfp;pdm2-Gal4/Lpp-Gal4* and *UAS-Fbp1*^*RNAi*^*/UAS-yki*^*S168A*^*-gfp;pdm2-Gal4/Lpp-Gal4*. Hexamerin density was calculated as the total signal intensity of each hexamerin divided by the tumor volume at each time point (x-axis, in h AED). For the 120h AED reference time point, corresponding to pupariation in *UAS-rfp*^*RNAi*^*/UAS-gfp;pdm2-Gal4/Lpp-Gal4* animals, hexamerin levels were measured in the wing disc pouch and normalized by pouch volume. 3-24 individual discs or tumors were analyzed per condition and time point.

To evaluate the functional importance of hexamerin uptake in supporting tumor growth, we performed volumetric reconstructions of tumors using confocal microscopy combined with 3D image analysis. In control tumor-bearing larvae, tumor volume increases rapidly between 144h and 216h AED, followed by a plateau that persists until larval death (Fig. 2B). In contrast, tumors from animals lacking *Fbp1* expression were significantly smaller at all time points starting from 168h AED (Fig. 2B), suggesting that albumin-like proteins uptake by the tumor is required for its expansion. Although an important part of hexamerins is found in the tumor at later stages (168-216h AED, compare Fig. 1E, F), it is possible that their reuptake in fat body cells followed by their breakdown into amino acids also contributes to tumor growth.

We then used tumor volume data to estimate the relative density of storage proteins within the tumor over time. We observed that intra-tumor hexamerin concentrations increased during the first 24-48h of the extended larval stage, after which they dropped markedly (Fig. 2C-F). This decline could reflect the combined effect of a progressive increase in tumor mass and the gradual depletion of circulating hexamerins as the fat body stops producing them.

Together, these findings indicate that FBP1-mediated uptake of albumin-like proteins provides nutrient to the tumor and that this process plays a critical role in fueling tumor growth.

### Cachectic and non-cachectic tumors reallocate hexamerins from the fat body to sustain their growth

Notably, the Yki^S168A^ tumor is known to induce a cachexia-like syndrome, characterized by systemic wasting of fat and muscle tissues(*9, 22*). Cachexia has been proposed to support tumor development by providing nutrients derived from host tissue breakdown(*9, 18, 23–25*). This raised the possibility that, in the context of cachexia, hexamerins might be redirected to the tumor due to impaired reuptake by the fat body.

To determine whether hexamerin uptake is specific to cachectic tumor models or a general feature of tumor development, we examined another type of tumors generated by silencing the *avalanche* gene(*26*). While these tumors originate in the same tissue as the *pdm2>yki*^*S168A*^*-gfp* model, their growth kinetics differs: after an initial period of moderate growth, *rn>avl*^*RNAi*^ tumors enter a phase of neoplastic expansion, causing a 2-3 day delay at pupariation and culminating in pupal lethality without inducing detectable cachexia(*10*). Immunofluorescence reveals that, similar to *pdm2>yki*^*S168A*^*-gfp* tumors, all hexamerins are present at high level in *rn>avl*^*RNAi*^ tumors compared to control wing discs (Fig. 3A, B). Gene expression confirmed that these proteins were not produced locally by the tumor (Fig. S5A). Moreover, hexamerin expression in the fat body was unchanged between control and tumor-bearing larvae (Fig. S5B), consistent with their production and secretion by the fat body followed by their uptake in the tumor. Using quantification of immunofluorescence signals, we observed that hexamerin levels increase steadily during the first 24-48h and subsequently declines (Fig. S5C-F), a profile similar to the one observed in the *pdm2>yki*^*S168A*^*-gfp* model. Silencing *Fbp1* resulted in a complete loss of hexamerins in tumors (Fig. S5C-F), demonstrating that FBP1-dependent import is a general feature of tumor-associated hexamerin uptake, rather than a consequence of fat body wasting caused by cachectic tumors.

**Fig. 3.**
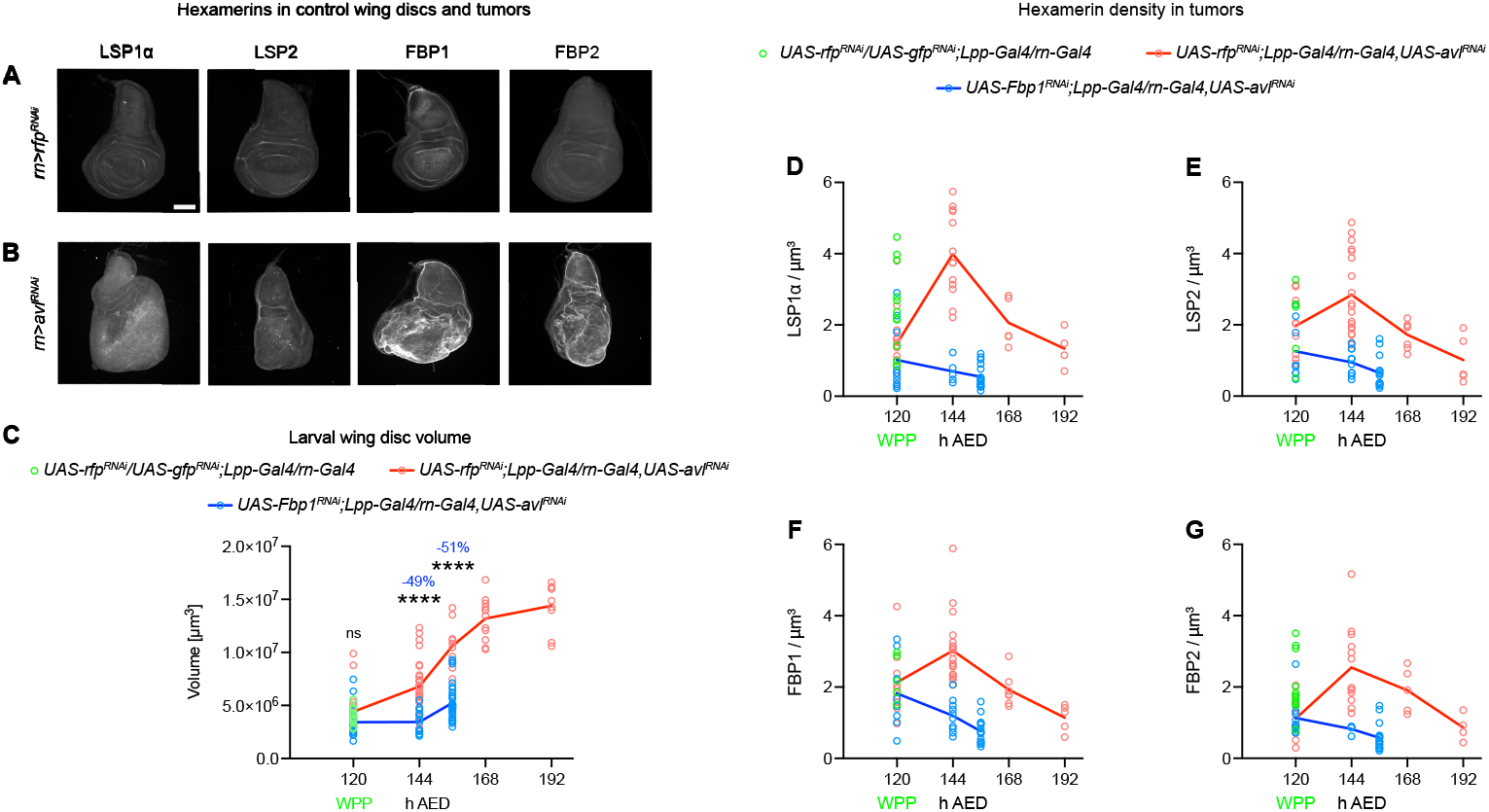
Hexamerins are taken up by *avl*^*RNAi*^ tumors through FBP1 action and contribute to tumor growth. (**A**) Representative images of wing imaginal discs dissected at 120h AED from control *rn>rfp*^*RNAi*^ white prepupae, stained with antibodies against LSP1α, LSP2, FBP1, and FBP2. Scale bar = 100 μm. (**B**) Representative images of tumors from wing discs of *rn>avl*^*RNAi*^ larvae, stained with the same antibodies and shown at the same magnification as in panel A. (**C**) Quantification of wing disc volume (y-axis) in tumor-bearing larvae expressing *avlRNAi*, either with FBP1 (*UAS-rfp*^*RNAi*^*;Lpp-Gal4/rn-Gal4,UAS-avl*^*RNAi*^) or with FBP1 knockdown (*UAS-Fbp1*^*RNAi*^*;Lpp-Gal4/rn-Gal4,UAS-avl*^*RNAi*^). Disc volumes were measured at the indicated time points (x-axis, in h AED). As a reference, control discs from *UAS-rfp*^*RNAi*^*/UAS-gfp*^*RNAi*^*;Lpp-Gal4/rn-Gal4* white prepupae were measured at 120h AED. For each genotype and time point, 9 to 33 wing discs were analyzed. Statistical significance was determined using unpaired *t*-tests or one-way ANOVA, as appropriate. ns = not significant; **** = p ≤ 0.0001. The percentage reduction in wing disc volume caused by FBP1 knockdown (*UAS-Fbp1*^*RNAi*^*;Lpp-Gal4/rn-Gal4,UAS-avl*^*RNAi*^) relative to *UAS-rfp*^*RNAi*^*;Lpp-Gal4/rn-Gal4,UAS-avl*^*RNAi*^ at 144h and 156h AED is indicated above the asterisks. (**D-G**) Time-course analysis of hexamerin density within tumors during the extended third instar stage. Intratumoral levels of LSP1α (**D**), LSP2 (**E**), FBP1 (**F**), and FBP2 (**G**) were measured in the following genotypes: *UAS-rfp*^*RNAi*^*;Lpp-Gal4/rn-Gal4,UAS-avl*^*RNAi*^ and *UAS-Fbp1*^*RNAi*^*;Lpp-Gal4/rn-Gal4,UAS-avl*^*RNAi*^. Hexamerin density was calculated by dividing the total signal intensity by tumor volume at each developmental time point (x-axis, in h AED). For reference, the 120h AED time point, corresponding to pupariation in *UAS-rfp*^*RNAi*^*/UAS-gfp*^*RNAi*^*;Lpp-Gal4/rn-Gal4* control animals, was used to determine baseline hexamerin density in wing discs. For each condition and time point, 4 to 19 individual discs or tumors were analyzed.

Analysis of tumor volume revealed that growth occurred during the period in which hexamerins accumulated (Fig. 3C). Importantly, when FBP1 was knocked down, tumor size was reduced by over 50% compared to controls (Fig. 3C), confirming that storage proteins are critical for supporting neoplastic growth.

Finally, we calculated hexamerin density within *rn>avl*^*RNAi*^ tumors by integrating protein accumulation data with volumetric measurements. This analysis showed that hexamerin density increased during the first 24h of the extended larval stage and declined sharply thereafter (Fig. 3D-G), likely reflecting the progressive depletion of systemic hexamerin stores and the continuous expansion of tumor mass. Together, these results establish that the uptake of albumin-like proteins both in cachectic (*pdm2>yki*^*S168A*^*-gfp*) and non-cachectic (*rn>avl*^*RNAi*^) tumors relies on a general FBP1-mediated mechanism that enables tumors to exploit host storage proteins as a source of nutrient.

### Hexamerin uptake by tumors modulates host developmental timing via *dilp8*

Both tumor models used in this study profoundly alter host development. While *pdm2>yki*^*S168A*^*-gfp* tumors cause a complete developmental arrest at the end of the larval stage (Fig. S1A), *rn>avl*^*RNAi*^ tumor-bearing larvae exhibit a delay of 2-3 days before pupariation(*10*).

Strikingly, upon *Fbp1* knockdown, the developmental arrest induced by *pdm2>yki*^*S168A*^*-gfp* tumors is largely rescued, pupariation occurring with only 11h delay compared to controls (Fig. 4A). Importantly, this rescue is specific to the loss of FBP1, as silencing other components of the protein storage system did not restore developmental progression (Fig. S6A).

**Fig. 4.**
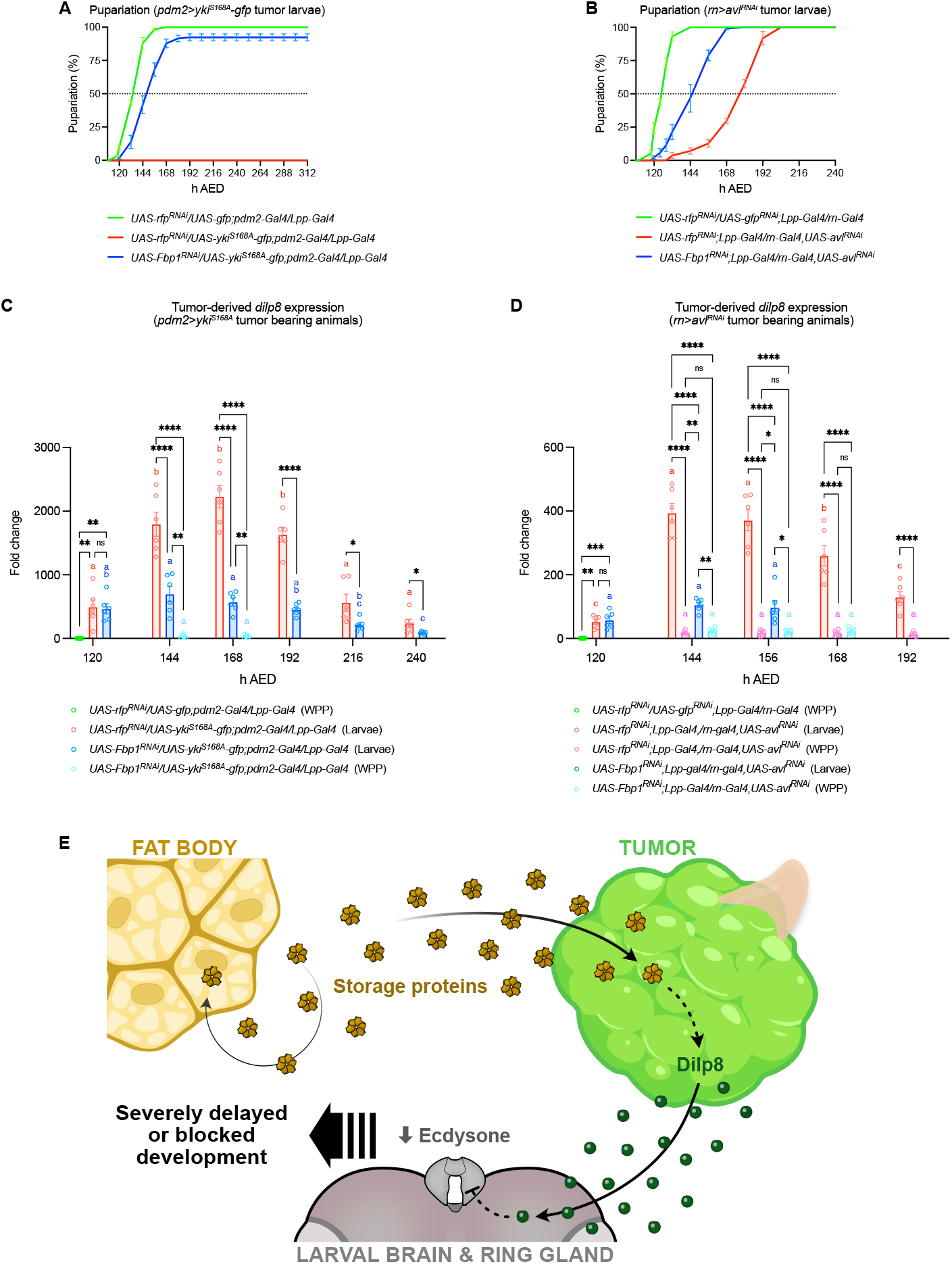
Hexamerins in tumors impair host development through enhanced *dilp8* expression. (**A, B**) Pupariation timing curves of larvae bearing *pdm2>yki*^*S168A*^*-gfp* (**A**) or *rn>avl*^*RNAi*^ (**B**) tumors in the presence (red lines) or absence (blue lines) of FBP1. Additional control genotypes are included (green lines). Time course data represent the mean ± SEM from three vials per genotype (N = 3). (**C**) Quantification of *dilp8* mRNA levels in dissected wing discs or tumors during the extended larval stage from the following genotypes: *UAS-rfp*^*RNAi*^*/UAS-gfp;pdm2-Gal4/Lpp-Gal4, UAS-rfp*^*RNAi*^*/UAS-yki*^*S168A*^*- gfp;pdm2-Gal4/Lpp-Gal4*, and *UAS-Fbp1*^*RNAi*^*/UAS-yki*^*S168A*^*-gfp;pdm2-Gal4/Lpp-Gal4*. Gene expression was measured by RT-qPCR. Bars show the mean ± SEM, with individual replicate values overlaid. Six biological replicates were analyzed per genotype at each time point (N = 6). Statistical differences *within each genotype* across time points were assessed using one-way ANOVA or unpaired *t*-tests, as appropriate, and are indicated by color-coded letters above the bars. Comparisons *between genotypes* at the same time point were also tested using one-way ANOVA or unpaired *t*-tests; significance is denoted by asterisks: ns = not significant, * = *p* ≤ 0.05, ** = *p* ≤ 0.01, **** = *p* ≤ 0.0001. Gene expression levels were normalized to *actin*. Developmental timing is reported in h AED. WPP: white prepupa. (**D**) RT-qPCR analysis of *dilp8* mRNA levels in *UAS-rfp*^*RNAi*^*/UAS-gfp*^*RNAi*^*;Lpp-Gal4/rn-Gal4* (control), *UAS-rfp*^*RNAi*^*;Lpp-Gal4/rn-Gal4,UAS-avl*^*RNAi*^, and *UAS-Fbp1*^*RNAi*^*;Lpp-Gal4/rn-Gal4,UAS-avl*^*RNAi*^. Expression data are shown as mean ± SEM with individual data points displayed. Six replicates were analyzed per condition, for each time point (N = 6). Statistical comparisons *within genotypes* across time were performed using one-way ANOVA or unpaired *t*-tests and represented by letters (same color) above bars. Comparisons *between genotypes* at the same time point were tested similarly and shown with asterisks: ns = not significant, * = *p* ≤ 0.05, ** = *p* ≤ 0.01, *** = *p* ≤ 0.001, **** = *p* ≤ 0.0001. Expression values were normalized to *actin*, and time points are given in h AED. WPP: white prepupa. (**E**) Model summarizing the interplay between tumor hexamerin uptake, *dilp8* expression, and systemic effects on host development. From the pool of storage proteins present in the hemolymph, only a small fraction is reabsorbed by the fat body, while the majority is taken up by the tumor via FBP1. This uptake induces the expression of Dilp8, which is secreted into the hemolymph and acts on the brain to reduce ecdysone synthesis in the prothoracic gland. The resulting lower ecdysone levels fail to trigger pupariation, leading to a severe delay or a complete block in host development.

Similarly, in the *rn>avl*^*RNAi*^ tumor model, blocking hexamerin import into the tumor partially alleviates the developmental delay, allowing earlier pupariation (Fig. 4B, Fig. S6B).

To investigate whether tumor size influences the ability to pupariate in this context, we measured tumor volumes in *Fbp1*-depleted, Yki^S168A^ tumor-bearing animals. We found that larvae that pupariated (white prepupae, WPP) showed only modestly, although significantly, smaller tumors than those developmentally arrested (Fig. S6C), suggesting that tumor size is not a major parameter contributing to the developmental delay.

Previous studies have shown that many types of tumorous imaginal discs secrete Dilp8, an insulin-like peptide that suppresses ecdysone production, thereby delaying metamorphosis(*10–14*). However, the dynamics of *dilp8* expression and its regulation by tumor metabolism remain elusive.

We therefore investigated *dilp8* transcription in relation to tumor growth and hexamerin uptake. In *pdm2>yki*^*S168A*^*-gfp* tumors, *dilp8* expression increases markedly at 120h AED and continues to rise to reach a plateau between 144h and 192h AED. After 192h AED, *dilp8* expression drops despite continued tumor growth (Fig. 4C, see also Fig. 2B). This pattern clearly indicates that *dilp8* levels do not scale linearly with tumor volume but rather follow an uncharacterized dynamic taking place in the tumor.

Interestingly, the *dilp8* plateau between 144h and 192h AED coincides with the timing of hexamerin accumulation in the tumor (see Fig. S4B-E). Upon *Fbp1* silencing, *dilp8* levels remained at very low levels in animals that pupariated (42% of the population at 144h AED and 88% at 168h AED, WPP in cyan on Fig. 4C) or reached intermediate levels in larvae that did not (58% of the population at 144h AED and 12% at 168h AED, Larvae in blue on Fig. 4C). This indicates that hexamerin uptake is a strong inducer of *dilp8* expression by the tumor and majorly contributes to delaying pupariation.

A similar pattern was observed in the *rn>avl*^*RNAi*^ tumor model: *dilp8* expression peaked between 144h and 156h AED, followed by a gradual decline despite continued tumor growth (Fig. 4D). As in the *pdm2>yki*^*S168A*^*-gfp* model, the rise in *dilp8* expression after 120h AED was abolished when hexamerin uptake was blocked by *Fbp1* knockdown (Fig. 4D).

Together, these results indicate that the production of Dilp8 does not scale with tumor volume but rather relies on a metabolic switch triggered by the uptake of hexamerins into tumor tissues. Therefore, tumors make use of hexamerin resources both to sustain their growth and to extend the larval period for prolonged tumor growth via Dilp8 production (Fig. 4E).

## DISCUSSION

A central challenge in tumor biology is to understand how proliferating cells meet their nutritional demands while concurrently manipulating systemic signaling for their benefit. In this study, we reveal a mechanism whereby epithelial tumors exploit circulating storage proteins both to fuel their growth and to delay host development.

Taking advantage of their role during *Drosophila* development(*8*), we show that hexamerins produced by the fat body are actively taken up by imaginal disc tumors during the larval stage. Their accumulation in tumors exhibits a biphasic pattern, with levels increasing during the first 24-48h of tumor growth followed by a decline. This phase of reduced accumulation likely results from both the depletion of circulating stores due to fat body exhaustion and intracellular degradation of internalized hexamerins into amino acids stores. Interestingly, the uptake of these storage proteins in the tumors requires the transporter FBP1, a hexamerin-binding protein previously implicated in regulating hexamerin uptake in fat body cells(*8*). This suggests that the same putative receptor might be used in both tissues for the binding of FBP1 and the subsequent receptor-mediated endocytosis of hexamerin complexes. In both cachectic and non-cachectic tumor models, loss of *Fbp1* impairs hexamerin uptake and significantly limits tumor growth. Therefore, hexamerin internalization is not a byproduct of systemic wasting but rather appears like a general feature promoting tumor growth.

Our findings echo earlier observations that mammalian serum albumins, structurally and functionally analogous to insect hexamerins, are taken up by tumor cells(*2, 3, 27*). While the mechanism of albumin uptake in human cancers remains incompletely characterized, it is increasingly clear that, particularly during the early stages of tumorigenesis, albumin does not enter tumor cells solely through passive mechanisms such as macropinocytosis or enhanced permeability and retention (EPR). Instead, it requires active import via specific receptors or chaperones, including SPARC and gp60 in glioblastoma and other cancers(*6, 28*). This is remarkably similar to what we observe in *Drosophila*, where hexamerin uptake by tumors relies on FBP1. The identification of FBP1 as a dedicated hexamerin for intracellular uptake in flies provides a genetically tractable platform to investigate how tumors co-opt protein-based nutrient stores and may serve to identify conserved pathways by which tumors exploit host macromolecular resources during early tumor progression.

We also uncover that beyond providing nutritional resources, hexamerin uptake in tumors also promotes systemic effects through the production of Dilp8, a hormone that delays pupariation. This delay provides tumors with additional time to expand before the onset of metamorphosis and the developmental feeding arrest. Blocking hexamerin uptake decreases *dilp8* expression and partially rescues developmental progression. Therefore, hexamerins coordinate internal amino acid stores and endocrine regulations.

This endocrine axis parallels the function of relaxins in mammals, a family of hormones related to Dilp8 and known to control tissue remodeling, reproduction, and metabolism(*29–31*). Intriguingly, relaxin-like peptides are often upregulated in human tumors where they contribute to tumor growth, matrix remodeling, and systemic metabolic changes such as cachexia(*32, 33*). Our findings suggest that the Dilp8/Relaxin pathway may represent a conserved module used by tumors to convert nutrient status into systemic hormonal signals that reprogram host physiology.

While the molecular cues that trigger *dilp8* induction upon hexamerin uptake remain unclear, our data support a model where tumors use hexamerin resources and other metabolic intermediates to activate hormonal signals and induce major physiological changes in the host. Deciphering these molecular cues will be an exciting focus for future research.

In summary, our study shows that tumors in *Drosophila* can act as systemic nutrient sinks, diverting host storage proteins to support their anabolic needs while simultaneously manipulating endocrine circuits to delay development. This dual implication of albumin-like proteins, as both metabolic substrates and regulators of systemic physiology, reveals a conserved evolutionary logic in tumor-host interactions.

These findings open new directions for exploring how tumors integrate nutrient sensing with hormonal control, and raise the possibility that targeting macromolecular nutrient scavenging, such as albumin uptake, may offer therapeutic leverage against tumor progression and associated metabolic syndromes.

## MATERIALS AND METHODS

### *Drosophila* strains and maintenance

Flies were maintained and all experiments conducted on a standardized diet consisting of 7.5g/L agar, 35g/L wheat flour, 50g/L yeast powder, 55g/L sugar, 25mL/L methyl, and 4 mL/L propionic acid. Experimental conditions were kept constant at 25°C. Both sexes were included in all assays. For precise developmental staging, eggs were collected over a 4h window on yeast-supplemented agar plates. L1 larvae were harvested 26h after the beginning of egg laying and transferred to vials containing standard food, with 40 larvae per vial. The exact developmental stage or time point used for each analysis is detailed in the relevant sections. The following stocks were obtained from the Bloomington Drosophila Stock Center (BDSC) at Indiana University: *pdm2-Gal4* (#49828), *UAS-gfp* (#39760), *Lpp-Gal4* (#84317), *rn-Gal4* (#7405), *UAS-rfp*^*RNAi*^ (#67852), and *UAS-gfp*^*RNAi*^ (#44412); while the following lines were obtained from the Vienna Drosophila Research Center (VDRC): *UAS-Lsp1*α^*RNAi*^ (#101101), *UAS-Lsp1γ*^*RNAi*^ (#38129), *UAS-Lsp2*^*RNAi*^ (#109979), *UAS-Fbp1*^*RNAi*^ (#330200), *UAS-Fbp2*^*RNAi*^ (#33172), and *UAS-avl*^*RNAi*^ (#107264). Other stocks used in this study were: *UAS-yki*^*S168A*^*-gfp*(*16*), and *rn-Gal4,UAS-avl*^*RNAi*^(*10*).

### Immunostainings and volume reconstruction

Wing discs and tumors were dissected from male and female larvae or white prepupae at the specified developmental stages in PBS. Tissues were fixed in 4% formaldehyde (ThermoFisher Scientific, #28908) for 30 minutes at room temperature, then washed in PBS with 0.3% Triton X-100 (PBT). After blocking in PBT containing 2% BSA, samples were incubated overnight at 4°C with primary antibodies. The following primary antibodies were used: rat anti-LSP1α(*8*) 1/200, rat anti-LSP2(*8*) 1/500, guinea pig anti-FBP1(*8*) 1/200, and guinea pig anti-FBP2(*8*) 1/1000.

On the following day, tissues were washed, re-blocked, and incubated for 2h at room temperature with secondary antibodies at 1/200 dilution. These included Goat anti-rat IgG (H+L) Highly Cross-Adsorbed Secondary Antibody, Alexa Fluor™ 647 (Invitrogen, A21247, Lot 1975526) and Goat anti-guinea pig IgG (H+L) Highly Cross-Adsorbed Secondary Antibody, Alexa Fluor™ 568 (Invitrogen, A11075, Lot 1970982).

Samples were mounted in Vectashield Antifade Medium with DAPI (Vector Laboratories, #H-1200-10) on glass-bottom Cellview dishes without coverslips to maintain native tissue architecture. Fluorescence imaging was performed on a Zeiss LSM900 Inverted Laser Scanning Confocal Microscope using Zen software (Zeiss). Full tissue volumes were captured by acquiring Z-stacks at 4.27μm intervals. Image processing and 3D surface reconstruction were carried out using Imaris software (Oxford Instruments)(*34*), and all conditions, including controls, were processed uniformly using Fiji.

### RT-qPCR

Larvae were collected at specific times after egg deposition (AED). Wing discs or tumors were dissected in cold PBS and immediately flash-frozen in liquid nitrogen. Total RNA was isolated using the RNeasy Plus Micro Kit (QIAGEN, #74034) following the manufacturer’s instructions. For each sample, 1μg of RNA was reverse-transcribed using a SuperScript IV VILO Master Mix (Invitrogen, #11756050). The resulting cDNA was used for quantitative PCR (qPCR) on a Viia 7 system (Applied Biosystems). Each cDNA sample was diluted 1:50 in water, then combined with 10μM gene-specific primers and *Power*SYBR Green PCR Master Mix (Applied Biosystems, #4367659). Gene expression was normalized to either *actin* or ribosomal protein *rp49* transcripts. Three independent biological replicates were analyzed, with each sample run in technical replicates.

Primers used included:

actin_For: TCGATCATGAAGTGCGACGT; actin_Rev: ACCGATCCAGACGGAGTACT.

rp49_For: CTTCATCCGCCACCAGTC; rp49_Rev: CGACGCACTCTGTTGTCG.

Lsp1α_For: GAGTACATTGCGATGGGAAAGC; Lsp1α_Rev: CATACGAGCGAAGGCCACAT.

Lsp1β_For: GATCGCCATCGCATTGCTG; Lsp1β_Rev: CCCTGCTTGATGTGGTCCT.

Lsp1*γ*_For: GCCTGTGTGACTGCCTTTAG; Lsp1*γ*_Rev: AGAGGCTCATCAATACGGTGA.

Lsp2_For: CTTCCAGCACGTCGTCTACTG; Lsp2_Rev: CCCTGCATATCATCACGGAACA.

Fbp1_For: ATCGTGGCGGCATTGATAAGG; Fbp1_Rev: CGAAGGGTGTCAAAGTCCTG.

Fbp2_For: ATGAATCTGACTGGCATGATCCA; Fbp2_Rev: CCAGGCCATAGACAGAGGACA.

dilp8_For: CGACAGAAGGTCCATCGAGT; dilp8_Rev: GATGCTTGTGCGTTTTG.

### Western blots

Wing discs, tumors, and fat bodies were collected from animals at defined developmental stages, then homogenized in ice-cold PBS supplemented with Halt™ Protease & Phosphatase Inhibitor Cocktail (ThermoFisher Scientific, #78440) by using a KIMBLE Dounce tissue grinder (Merck, #D8938). Homogenates were immediately snap-frozen in liquid nitrogen. For hemolymph collection, larvae were bled into a drop of ice-cold PBS containing the same inhibitor cocktail. Serum was clarified by repeated centrifugation at 4°C to eliminate cellular contaminants. Protein extracts were prepared from the same number of animals or dissected organs collected at different timepoints. Samples were mixed with 4xLaemmli Sample Buffer (Bio-Rad, #1610747) containing 10% 2-Mercaptoethanol (Sigma-Aldrich, #M6250), then boiled at 95°C for 10 minutes. Proteins were separated by SDS-PAGE using Mini-PROTEAN TGX Precast Gels (Bio-Rad, #4569033) in Tris/Glycine/SDS Buffer (Bio-Rad, #1610732) and transferred to 0.2μm nitrocellulose membranes with the Trans-Blot Turbo system (Bio-Rad, #1704158). Membranes were blocked in Tris buffered saline (Bio-Rad, #1706435) + 0.1% Tween-20 + 5% BSA, then incubated with the following primary antibodies: LSP1α (1:1000)(*8*), LSP2 (1:5000)(*8*), FBP1 (1:1000)(*8*), and FBP2 (1:5000)(*8*). Loading controls included Actin (Sigma-Aldrich, #A2103; 1:5000) and Cv-d (1:1000)(*21*). Horseradish peroxidase (HRP)–conjugated secondary antibodies (Invitrogen, #31470; Jackson ImmunoResearch, #111-035-144; Invitrogen, #A18769; 1:5000) were used with the Clarity Max™ Western ECL Substrate (Bio-Rad, #1705062) for signal detection. All results are representative of three independent biological replicates. Band intensities were quantified using Fiji software.

### Hexamerin absolute signal quantification and its density

After staining procedures were completed, tissues were placed in glass-bottom Cellview dishes containing Vectashield Antifade Mounting Medium with DAPI, omitting the use of coverslips. Imaging was carried out using a Zeiss LSM900 inverted confocal microscope operated via Zen software (Zeiss). To reconstruct entire tissue volumes, Z-stack images were collected at 4.27μm intervals.

To quantify the total hexamerin signal within tumors, we developed a custom Fiji macro (code in Fig. S2). First, images were converted to TIFF format. Tumor regions were segmented using either GFP fluorescence or DAPI staining, depending on the genotype. Specifically, for *pdm2>yki*^*S168A*^*-gfp* tumors, GFP fluorescence was used as a mask to delineate the tumor area. In contrast, for *rn>avl*^*RNAi*^ tumors, which lacked GFP labeling, DAPI staining was used to define the tumor boundaries. Within each Z-slice, the macro measured (i) the mean gray value of the hexamerin signal within the segmented tumor region, and (ii) the area of this region. Since the mean gray value is the sum of the gray values of all the pixels in the selection divided by the number of pixels, multiplying (i) by (ii) yielded the absolute hexamerin signal for that slice. The total signal for the tumor was obtained by summing the absolute values across all slices. This cumulative signal was subsequently normalized to the tissue volume (in μm^3^) to compute the hexamerin signal density.

### Food intake

Food intake was measured using a dye-based assay, as previously reported(*35*). Synchronized third-instar larvae were rinsed in PBS, blotted dry with a Kimwipe, and placed on standard fly food containing 1.5% (w/v) blue dye (Erioglaucine Disodium Salt, Merck, #861146) for 1h at 25°C. After feeding, larvae were washed in PBS, dried, weighed, flash-frozen in liquid nitrogen, and stored at -20°C. For analysis, larvae were homogenized in 20µl PBS and spun down at 4°C for 20 minutes. A 10µl sample of the supernatant was transferred to a clean tube, and dye levels were quantified by measuring absorbance at 629nm using a spectrophotometer. Results were then normalized to the corresponding larval mass.

### Wing imaginal disc volume

Wing discs were dissected from larvae at 120h AED in ice-cold PBS, then fixed in 4% formaldehyde for 20 minutes. Tissues were mounted in Vectashield Antifade Medium with DAPI on Cellview glass-bottom culture dishes without coverslips. Imaging was carried out using a Zeiss LSM900 inverted confocal microscope and Z-stack images were processed with Imaris software(*34*).

### Growth curves

Adult females laid eggs for 4h on plates containing PBS, 2% agar, and 2% glucose. After 26h, forty first-instar larvae were transferred to tubes with standard fly food and kept at 25°C. To track pupariation timing, larvae were monitored every 2-3h, and the number entering pupariation was recorded.

For eclosion timing, five groups of forty synchronized white prepupae per genotype were followed, and newly emerged adults were counted to generate eclosion curves.

### Adult wing area

Adult females of the desired genotype were collected and preserved in ethanol. Wings were dissected and mounted in a 6:5 lactic acid to ethanol solution. Images were captured at 1024×768 resolution using a Leica MZ16-FA fluorescence stereomicroscope equipped with a DFC-490 digital camera in bright-field mode (settings: 50% light intensity, 10.5 exposure, 2.3 gain, 152 saturation, 1.2 gamma). Wing area was quantified using a previously established deep learning-based segmentation method(*34*).

## SUPPLEMENTARY MATERIALS

The PDF includes:

Figs. S1 to S6

## ACKNOWLEDGMENTS

We thank Allison Bardin, Laura Boulan and members of the laboratory for discussions and comments on the manuscript; the Bloomington Stock Center and the Vienna *Drosophila* Resource Center for providing fly stocks; the PICT-IBiSA@BDD light-microscopy facility of Institut Curie. This work was supported by Institut Curie, CNRS, INSERM, FRM, European Research Council (Advanced Grant n°694677 to P.L.), Labex DEEP program (ANR-11-LABX-0044, ANR-10-IDEX-0001-02).

## AUTHOR CONTRIBUTIONS

Conceptualization, L.V. and P.L.; Methodology, L.V., D.B.-O., A.A. and P.L.; Investigation, L.V.; Writing – Original Draft, L.V.; Writing – Review & Editing, L.V. and P.L.; Funding Acquisition, P.L.; Supervision, L.V. and P.L.

