## Supplementary Materials for "*Drosophila* storage proteins promote both the rate and the duration of tumor growth"

### SUPPLEMENTARY MATERIALS
### (FIGURES S1-S6 AND CORRESPONDING LEGENDS)

#### Figure S1

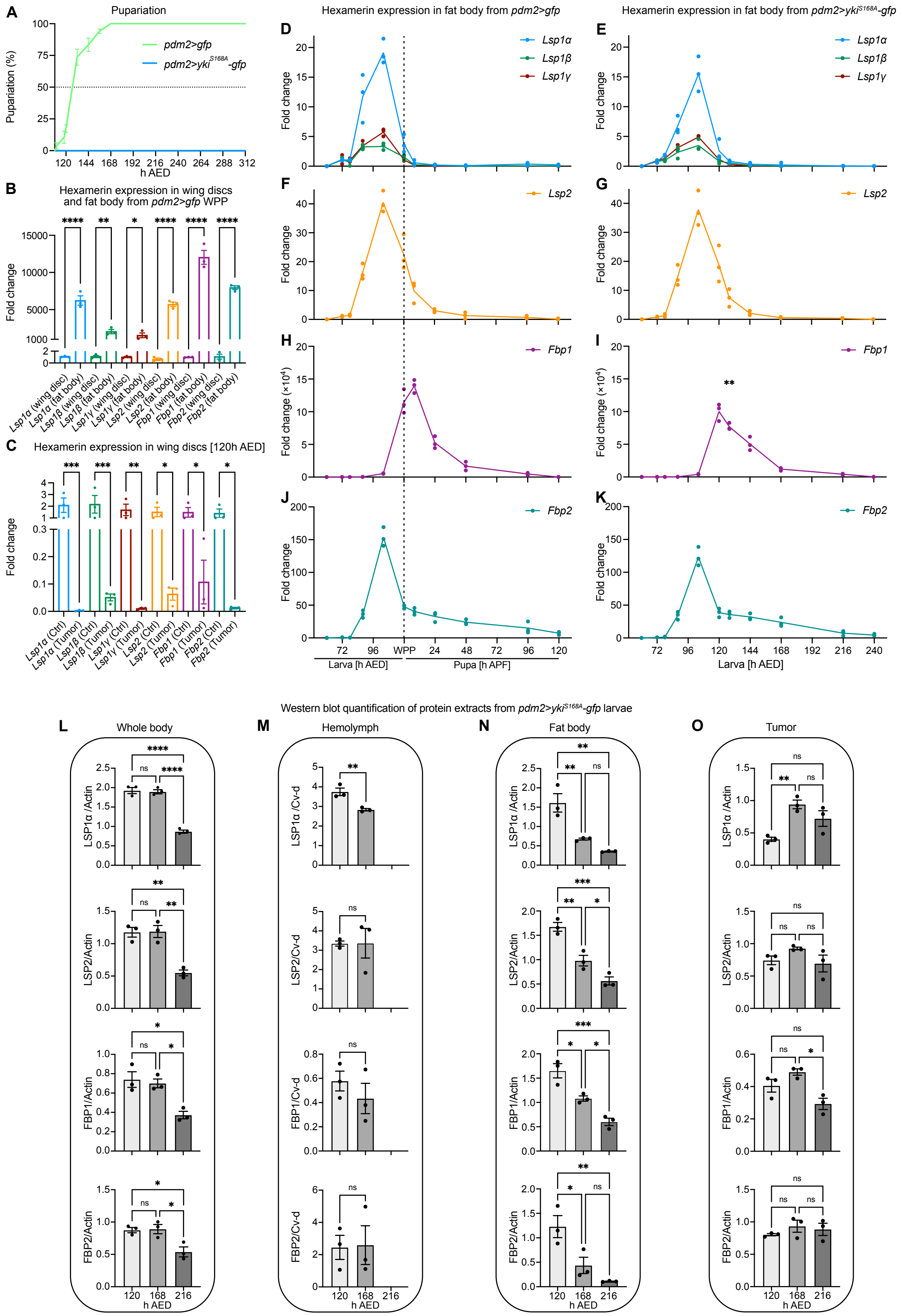

**Fig. S1**. **Yki$^{S168A}$ tumors scavenge circulating hexamerins without inducing their expression or producing them locally.** (**A**) Timing of pupariation for *pdm2>gfp* and *pdm2>yki$^{S168A}$-gfp* animals. Mean values ± SEM are plotted over time (x-axis: h AED). For each genotype, N = 3 represents the mean from three separate vials in a single experiment. (**B**) RT-qPCR analysis of *Lsp* and *Fbp* transcripts in wing discs and fat bodies isolated from *pdm2>gfp* white prepupae (WPP). Each data point corresponds to an individual replicate; bars indicate mean ± SEM. Significance was evaluated by unpaired t test. * = $p \leq 0.05$; ** = $p \leq 0.01$; **** = $p \leq 0.0001$. N = 3 replicates per gene. Expression was normalized to *rp49*. (**C**) Transcript levels of *Lsp* and *Fbp* genes in wing discs from *pdm2>gfp* (Ctrl) and *pdm2>yki$^{S168A}$-gfp* (Tumor) animals at 120h AED, measured by RT-qPCR. Each dot represents an independent replicate; bars show mean ± SEM. Statistical significance was assessed using an unpaired t test. * = $p \leq 0.05$; ** = $p \leq 0.01$; *** = $p \leq 0.001$. N = 3 replicates per gene. Expression levels were normalized to *rp49*. (**D-K**) Developmental expression profiles of *Lsp1*$\alpha$, *Lsp1*$\beta$, *Lsp1*$\gamma$ (**D**, **E**), *Lsp2* (**F**, **G**), *Fbp1* (**H**, **I**), and *Fbp2* (**J**, **K**) in control (*pdm2>gfp*, left panels) and tumor-bearing (*pdm2>yki$^{S168A}$-gfp*, right panels) animals, analyzed by RT-qPCR at indicated developmental time points. Dotted lines mark the timing of the larva-to-pupa transition in *pdm2>gfp* animals and the equivalent time point in *pdm2>yki$^{S168A}$-gfp* animals, which fail to undergo pupariation. Each dot represents a replicate; lines connect mean values across time. Comparisons at each time point between genotypes were performed using Welch's t test. ** = $p \leq 0.01$. N = 3 replicates per genotype for each time point. Time is shown as hours after egg deposition (h AED) and hours after puparium formation (h APF). All expression levels were normalized to *rp49*. (**L-O**) Quantification of LSP1α, LSP2, FBP1, and FBP2 protein levels in whole-body lysates (**L**), hemolymph (**M**), fat body (**N**), and tumor (**O**) tissues of *pdm2>yki$^{S168A}$-gfp* animals at 120, 168, and 216h AED. Protein levels were normalized to Cv-d in hemolymph samples, and to actin in whole body, fat body, and tumor samples. Data are shown as individual replicates with mean ± SEM. Statistical significance was determined using Tukey's multiple comparison test. * = $p \leq 0.05$; ** = $p \leq 0.01$; *** = $p \leq 0.001$; **** = $p \leq 0.0001$. N = 3 replicates per time point. Corresponding western blots are displayed in Fig. 1C-F.

**Figure S2**

```
// Define the macro
macro "Process Images in Directory" {
    run("Set Measurements...", "area mean display redirect=None decimal=3");

    // Ask the user to select a directory
    dir = getDirectory("Choose a Directory");

    // Get all files in the directory
    fileList = getFileList(dir);

    // Loop through each file in the directory
    for (i = 0; i < fileList.length; i++) {
        filename = fileList[i];
        fullPath = dir + filename;

        // Only process .tif images
        if (endsWith(filename, ".tif") || endsWith(filename, ".TIF")) {
            print("Processing: " + filename);
            open(fullPath);
            // Get the number of channels in the active image
            Stack.getDimensions(width, height, channels, slices, frames);
            if (channels == 4) {
                run("Arrange Channels...", "new=124");
            } else {
            print("Image has " + channels + " channels. No changes made.");
            }
            run("Duplicate...", "duplicate channels=1-3");
            run("8-bit");
            title = getTitle();
            run("Split Channels");

            // Loop through each slice
            for (slice = 1; slice <= nSlices; slice++) {


                // Apply threshold to detect dapi signal (adjust as needed)
                selectWindow("C3-" + title);
                run("Gaussian Blur...", "sigma=2 stack");
                run("Auto Threshold", "method=Li ignore_black white stack");
                setSlice(slice);
                run("Create Selection");
                selectWindow("C1-" + title);
                setSlice(slice);
                run("Restore Selection");
                run("Measure");
                selectWindow("C2-" + title);
                setSlice(slice);
                run("Restore Selection");
                run("Measure");

            }

            run("Close All"); // Close the image after processing
        }
    }
    selectWindow("Results");
    saveAs("Results", dir + "results.csv");
}
```

**Fig. S2**. **Fiji macro for quantifying total hexamerin signal in tumors.** Code of the custom Fiji macro used to extract hexamerin signal measurements from tumor tissues. Confocal Z-stacks (4.27 µm intervals) of stained samples were converted to TIFF format, and tumor regions were segmented using either GFP fluorescence (*pdm2>yki*$^{S168A}$*-gfp* tumors) or DAPI staining (*rn>avl*$^{RNAi}$ tumors without GFP). For each Z-slice, the macro outputs (i) the mean gray value of the hexamerin staining within the segmented tumor area and (ii) the corresponding area. These two values were then multiplied by the operator to obtain the absolute hexamerin signal per slice. Summing these absolute values across all slices yielded the total hexamerin signal for the tumor, which was subsequently normalized to the tissue volume (µm$^3$) to calculate the hexamerin signal density.

### Figure S3

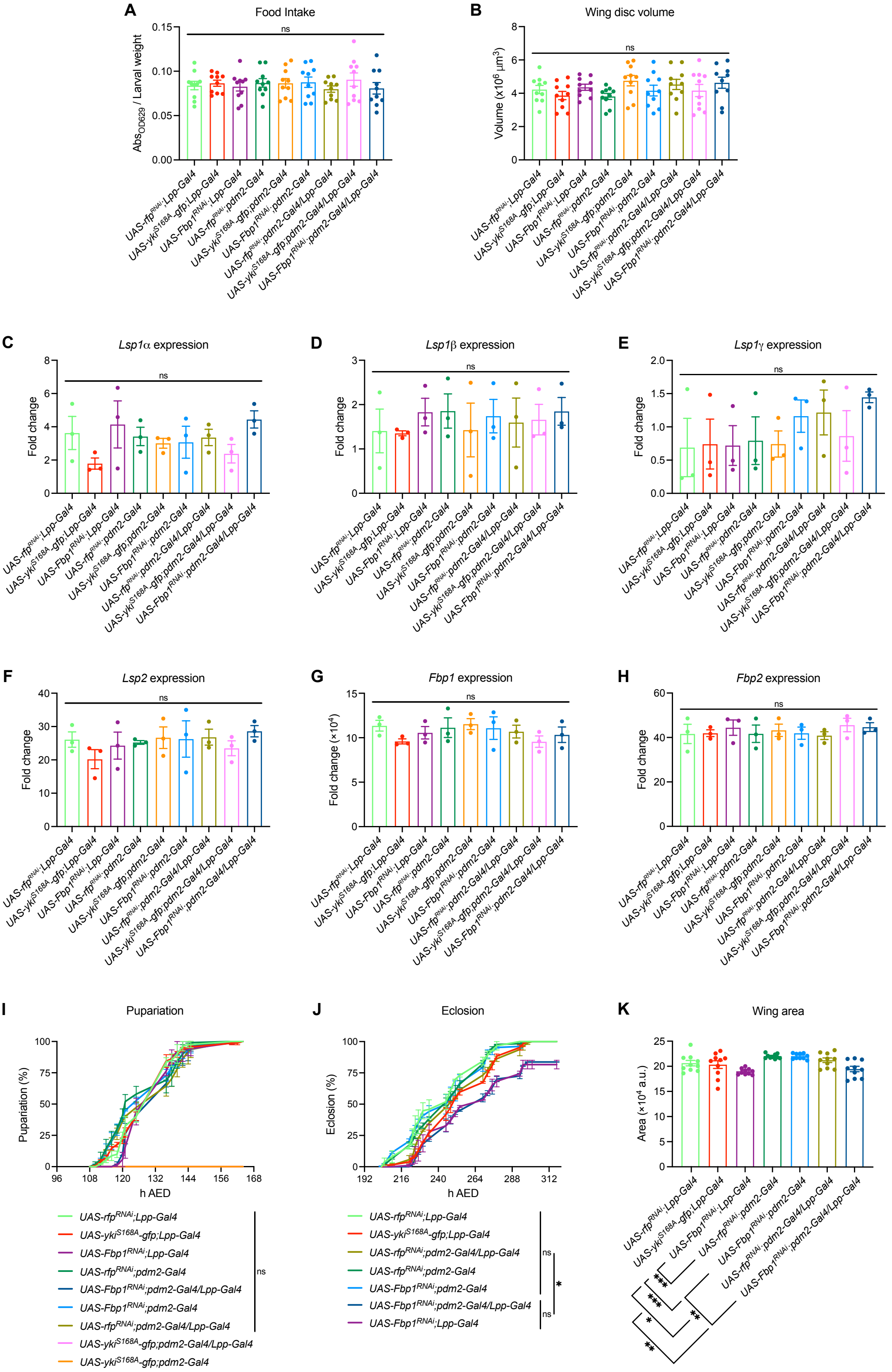
A
Food Intake
$Abs_{OD629}$ / Larval weight
ns
B
Wing disc volume
Volume (x$10^6$ µm$^3$)
ns
C
Lsp1α expression
Fold change
D
Lsp1β expression
E
Lsp1γ expression
F
Lsp2 expression
G
Fbp1 expression
Fold change (×$10^4$)
H
Fbp2 expression
I
Pupariation
Pupariation (%)
h AED
J
Eclosion
Eclosion (%)
K
Wing area
Area (×$10^4$ a.u.)
UAS-rfp$^{RNAi}$;Lpp-Gal4
UAS-yki$^{S168A}$-gfp;Lpp-Gal4
UAS-Fbp1$^{RNAi}$;Lpp-Gal4
UAS-rfp$^{RNAi}$;pdm2-Gal4
UAS-yki$^{S168A}$-gfp;pdm2-Gal4
UAS-Fbp1$^{RNAi}$;pdm2-Gal4
UAS-rfp$^{RNAi}$;pdm2-Gal4/Lpp-Gal4
UAS-yki$^{S168A}$-gfp;pdm2-Gal4/Lpp-Gal4
UAS-Fbp1$^{RNAi}$;pdm2-Gal4/Lpp-Gal4

**Fig. S3. Using two Gal4/UAS systems to simultaneously induce wing disc tumors and knock down *Fbp1* do not produce confounding developmental side effects.** (**A**) Larval feeding behavior was assessed by measuring ingestion of Erioglaucine blue dye-supplemented food, normalized to body weight. Data were analyzed using Šidák's multiple comparisons test. No statistically significant differences were observed. N = 10 larvae per genotype. (**B**) Wing disc size was measured in larvae of the indicated genotypes. Individual data points represent disc volumes, with mean ± SEM shown. Statistical comparisons were performed using Šidák's multiple comparisons test. No significant differences were detected. N = 10 wing discs per genotype. (**C-H**) Relative expression levels of hexamerin genes were quantified by RT-qPCR at 120h AED in larvae of the indicated genotypes. All expression levels were normalized to *rp49.* Values represent mean ± SEM. Statistical significance was determined by Tukey's multiple comparisons test. ns = not significant. N = 3 biological replicates per genotype. (**I**) Pupariation timing was tracked in larvae of each genotype, with mean ± SEM across time points plotted as connected lines. Analysis was conducted via two-way ANOVA with multiple comparisons. No significant differences were found. N = 3, representing the average from three independent vials per genotype. (**J**) Adult eclosion timing was monitored and is presented as mean ± SEM over time, with comparisons made by two-way ANOVA multiple comparisons test. N = 3, each representing the mean from three replicate vials per genotype. ns = not significant; * = $p \leq 0.05$. (**K**) Adult wing area was measured in females of the indicated genotypes. Mean ± SEM is shown, and significant differences were assessed using Šidák's multiple comparisons test. Only significant results are reported: * = $p \leq 0.05$; ** = $p \leq 0.01$; *** = $p \leq 0.001$. N = 10 wings per genotype.

**Figure S4**

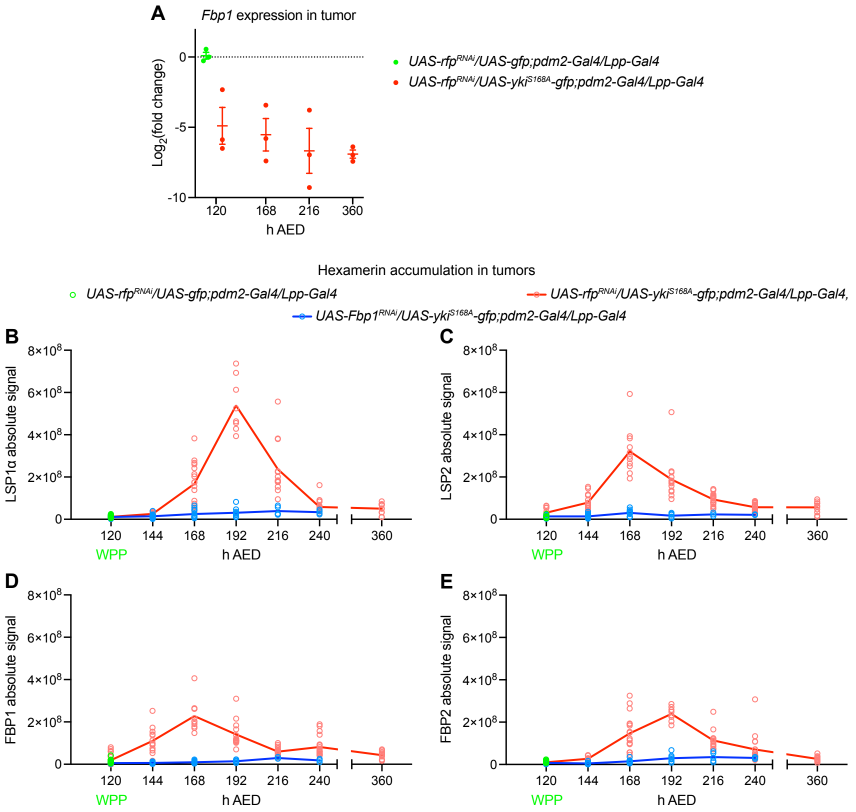


**Fig. S4. Yki$^{S168A}$ tumor progression is marked by dynamic changes in hexamerin levels while *Fbp1* tumor expression remains minimal.** (**A**) Time-course analysis of *Fbp1* expression in wing discs from larvae of the indicated genotypes, measured at multiple developmental stages (x-axis, in hours after egg deposition). Expression values are presented as $Log_2$(fold change), normalized to *rp49*. Each point represents an individual replicate (N = 3 per time point and genotype), with mean ± SEM shown. (**B-E**) Quantification by immunostaining of absolute signal intensity for LSP1α (**B**), LSP2 (**C**), FBP1 (**D**), and FBP2 (**E**) within tumors during the extended last instar larval stage in animals developing tumors either in the presence (*UAS-rfp$^{RNAi}$/UAS-yki$^{S168A}$-gfp;pdm2-Gal4/Lpp-Gal4*) or absence (*UAS-Fbp1$^{RNAi}$/UAS-yki$^{S168A}$-gfp;pdm2-Gal4/Lpp-Gal4*) of FBP1. Time points (in h AED) are indicated on the x-axis. The 120h AED time point corresponds to pupariation in the control genotype (*UAS-rfp$^{RNAi}$/UAS-gfp;pdm2-Gal4/Lpp-Gal4*), in which hexamerins were quantified in the wing disc pouch. For each time point, 4 to 18 wing discs/tumors were analyzed per condition.

**Figure S5**

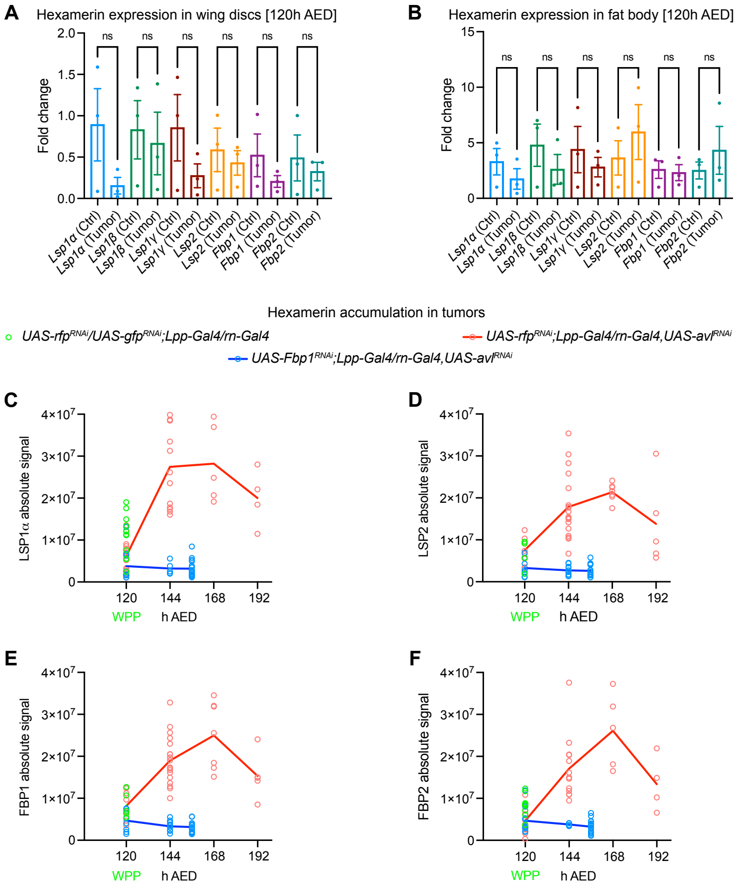


**Fig. S5. *avl*$^{RNAi}$ tumor does not alter hexamerin expression yet promotes their intratumoral accumulation.** (**A**) Relative expression of *Lsp* and *Fbp* genes in wing discs from *rn>rfp*$^{RNAi}$ (Ctrl) and *rn>avl*$^{RNAi}$ (Tumor) animals at 120h (AED), measured by RT-qPCR. Each dot represents a biological replicate (N = 3 per gene); bars show mean ± SEM. Data were normalized to *rp49*, and statistical comparisons were performed using unpaired t tests. No significant differences were observed (ns). (**B**) RT-qPCR quantification of *Lsp* and *Fbp* transcripts in fat bodies from *rn>rfp*$^{RNAi}$ (Ctrl) and *rn>avl*$^{RNAi}$ (Tumor) animals at 120h AED. Individual replicates (N = 3 per gene) are shown, with mean ± SEM. Gene expression was normalized to *rp49*, and unpaired t tests indicated no significant changes (ns). (**C-F**) Quantification of absolute signal intensity for LSP1α (**C**), LSP2 (**D**), FBP1 (**E**), and FBP2 (**F**) within tumors through immunofluorescence during the extended last instar larval stage. Tumors were analyzed in animals expressing *avl*$^{RNAi}$ either in the presence (*UAS-rfp*$^{RNAi}$*;Lpp-Gal4/rn-Gal4,UAS-avl*$^{RNAi}$) or absence (*UAS-Fbp1*$^{RNAi}$*;Lpp-Gal4/rn-Gal4,UAS-avl*$^{RNAi}$) of FBP1. Time points are indicated on the x-axis (in h AED). The 120h AED time point corresponds to pupariation of control animals (*UAS-rfp*$^{RNAi}$*/UAS-gfp*$^{RNAi}$*;Lpp-Gal4/rn-Gal4*), in which hexamerin levels were assessed in the wing discs. Between 4 and 19 wing discs or tumors were analyzed per genotype at each time point.

**Figure S6**

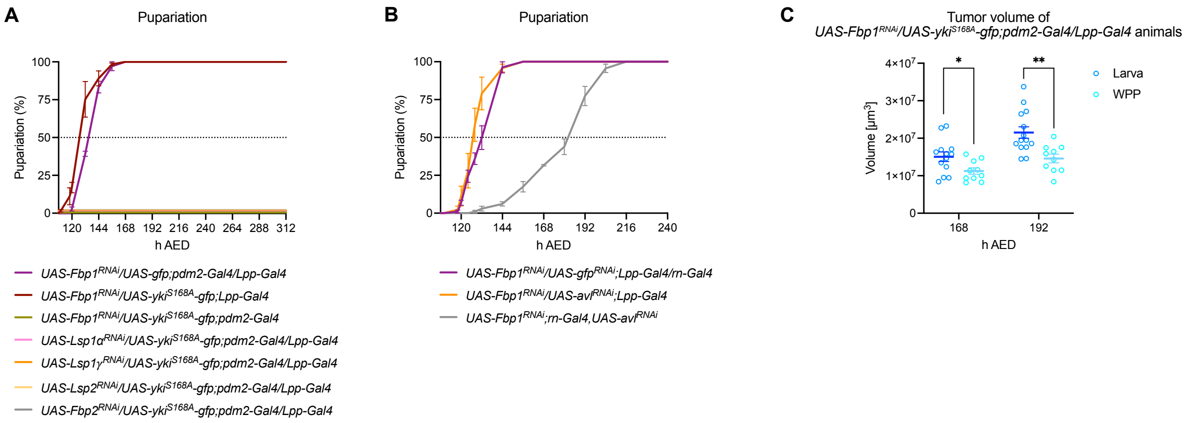


**Fig. S6. Hexamerin uptake by tumors and its contribution to tumor growth impact host developmental progression.** (**A**) Timing of pupariation in larvae of the indicated genotypes. These data are derived from the same experimental cohort shown in Fig. 4A. Line graphs represent the mean pupariation percentages ± SEM at various time points (x-axis: h AED). For each genotype, N = 3 corresponds to the average of three independent vials from a single experiment. (**B**) Pupariation curves for the indicated genotypes, based on the same dataset shown in Fig. 4B. Mean values ± SEM are plotted over time (x-axis: AED, in hours). For each genotype, N = 3 represents the mean from three separate vials in a single experiment. (**C**) Quantification of tumor volume in *UAS-Fbp1$^{RNAi}$/UAS-yki$^{S168A}$-gfp;pdm2-Gal4/Lpp-Gal4* larvae (blue) and white prepupae (WPP, cyan) at 168h and 192h AED. Tumor sizes were compared between larval and white prepupal stages using unpaired t-tests at each time point. * = $p \leq 0.05$; ** = $p \leq 0.01$. For each time point, a total of 10 to 14 tumors were measured per condition.
